# The vestibulospinal nucleus is a locus of balance development

**DOI:** 10.1101/2023.12.06.570482

**Authors:** Kyla R. Hamling, Katherine Harmon, Yukiko Kimura, Shin-ichi Higashijima, David Schoppik

## Abstract

Mature vertebrates maintain posture using vestibulospinal neurons that transform sensed instability into reflexive commands to spinal motor circuits. Postural stability improves across development. However, due to the complexity of terrestrial locomotion, vestibulospinal contributions to postural refinement in early life remain unexplored. Here we leveraged the relative simplicity of underwater locomotion to quantify the postural consequences of losing vestibulospinal neurons during development in larval zebrafish of undifferentiated sex. By comparing posture at two timepoints, we discovered that later lesions of vestibulospinal neurons led to greater instability. Analysis of thousands of individual swim bouts revealed that lesions disrupted movement timing and corrective reflexes without impacting swim kinematics, and that this effect was particularly strong in older larvae. Using a generative model of swimming, we showed how these disruptions could account for the increased postural variability at both timepoints. Finally, late lesions disrupted the fin/trunk coordination observed in older larvae, linking vestibulospinal neurons to postural control schemes used to navigate in depth. Since later lesions were considerably more disruptive to postural stability, we conclude that vestibulospinal contributions to balance increase as larvae mature. Vestibulospinal neurons are highly conserved across vertebrates; we therefore propose that they are a substrate for developmental improvements to postural control.

**SIGNIFICANCE STATEMENT:** Many animals experience balance improvements during early life. Mature vertebrates use vestibulospinal neurons to transform sensed instability into postural corrections. To understand if/how these neurons shape postural development, we ablated them at two developmentally important timepoints in larval zebrafish. Loss of vestibulospinal neurons disrupted specific stabilizing behaviors (swim timing, tilt correction, and fin/body coordination) more profoundly in older fish. We conclude that postural development happens in part by changes to vestibulospinal neurons — a significant step towards understanding how developing brains gain the ability to balance.

## INTRODUCTION

Animals actively modulate the timing and strength of their trunk and limb movements to remain balanced during locomotion^1–5^. In many species, balance control improves during early postnatal life as the behavioral strategies to correct imbalance refine. Understanding these sensorimotor computations, and how they refine over time, requires identifying the neurons that contribute towards balance development. Vestibulospinal neurons are descending projection neurons conserved across vertebrates that are well-poised to regulate balance^6–12^. They have somata in the lateral vestibular nucleus^13,14^ and receive convergent excitatory^15,16^ and inhibitory vestibular input^17,18^ as well as a wide variety of extravestibular inputs^19^.

The complexity of the mammalian nervous system and of tetrapod locomotion constrain our understanding of the contribution of vestibulospinal neurons to behavior. In decerebrate cats, vestibulospinal neurons encode body tilts, are active during extensor muscle contraction during locomotion^16,20^–22, and relay VIII^th^ nerve activity to ipsilateral extensor muscles^23,24^. Targeted loss-of-function experiments established the necessity of vestibulospinal neurons for specific hindlimb extension reflexes following imposed instability^25,26^. Questions remain as to what role vestibulospinal neurons play in balance control during natural movements and whether their role changes as balance matures.

Swimming offers a tractable means to assay neuronal contributions to balance computations across development. The biophysical and biomechanical contributions to swimming are straightforward to delineate^27,28^. For example, larval zebrafish swim with short discontinuous propulsive movements called “bouts” that constitute active locomotion. Between bouts, they are passive, akin to standing still. This dichotomy allows for dissociation of passive and active (i.e. neuronal) contributions to stability. Larval zebrafish maintain their preferred near-horizontal pitch using two active computations: they initiate swim bouts to occur when pitched off-balance and they rotate their bodies during bouts to restore posture^29–31^. Similar corrections stabilize posture in the roll axis^32^. Importantly, both behaviors improve over the first week of life^29–31^.

Vestibulospinal neurons might subserve vertebrate postural development. In mammals, vestibulospinal pathways have functional synapses at birth^33^. Teleost vestibulospinal neurons share anatomy^11,34^,35, ontogeny^36^, and functional responses^37,38^ with mammals. In larval zebrafish, vestibulospinal neurons encode vestibular stimuli as early as 4 days post-fertilization (dpf)^35,37^–40, Further, loss of the vestibular periphery disrupts trunk/fin coordination that larval zebrafish use during early life to climb in depth^41^.

Here, we defined the contribution of vestibulospinal neurons to postural orientation in freely swimming fish during early development. We selected two important timepoints: 4 dpf, when fish begin to orient and swim freely, and 7 dpf, after key stabilizing computations have matured. After targeted lesions, fish could still swim and continued to orient and navigate in depth. Posture was more variable after lesions at both timepoints; this effect was stronger at 7 than at 4 dpf. Further, lesions disrupted posture-dependent bout initiation and restorative rotations with stronger effects at 7 dpf. A model of swimming could account for postural variability following lesions at both ages. Finally, fin-body coordination was impaired after vestibulospinal loss at 7, but not 4 dpf. Altogether, our work reveals a specific contribution of vestibulospinal neurons to postural stability and locomotion across early development. Given the near-ubiquity of vestibulospinal neurons across vertebrates, and the shared challenge of postural development, we propose that this circuit may serve as a substrate for developmental improvements to balance.

## MATERIALS AND METHODS

### Fish Care

All procedures involving zebrafish larvae (*Danio rerio*) were approved by the Institutional Animal Care and Use Committee of New York University. Fertilized eggs were collected and maintained at 28.5°C on a standard 14/10 hour light/dark cycle. All experiments were performed on larvae between 4 and 9 dpf. During this time, zebrafish larvae have not yet differentiated their sex into male/female. Before 5 dpf, larvae were maintained at densities of 20-50 larvae per petri dish of 10 cm diameter, filled with 25-40 mL E3 with 0.5 ppm methylene blue. After 5 dpf, larvae were maintained at densities under 20 larvae per petri dish and were fed cultured rotifers (Reed Mariculture) daily.

### Fish Lines

Experiments were done on the mitfa-/-background to remove pigment. Larvae for vestibulospinal lesions were labeled with the double transgenic *Tg(nefma:hsp70l-LOXP-GAL4ff)*;*Tg(UAS:EGFP)*, henceforth called *Tg(nefma::EGFP)*^37^. Photoconverted larvae used for additional vestibulospinal lesions were from the *Tg(α-tubulin:C3PA-GFP)*^42^ background.

### Labeling Vestibulospinal Neurons with Spinal Photoconversions

PA-GFP positive larvae were raised in a dark incubator to prevent background photoconversion. Larvae were anesthetized with 0.2 mg/mL ethyl-3-aminobenzoic acid ethyl ester (MESAB, Sigma-Aldrich E10521, St. Louis, MO) and mounted laterally in 2% low-melting temperature agarose (Thermo Fisher Scientific 16520). Using a Zeiss LSM800 confocal microscope with 20x objective (Zeiss W Plan-Apochromat 20x/1.0 DIC CG=0.17 M27 75mm), the spinal cord between the mid-point of the swim bladder and the caudal-most tip of the tail was repeatedly scanned with a 405 nm laser until fully converted. For retrograde labelling of vestibulospinal neurons used for photoablations, the spinal cords of *Tg(α-tubulin:C3PA-GFP)* larvae were converted at 6 dpf. To allow the converted fluorophore to diffuse into neuron bodies, all fish were removed from agarose after photoconversion and raised in E3 in a dark incubator for 18-24 hours before imaging. Using this method, all spinal-projecting neurons are labeled with photoconverted PA-GFP.

### Spinal Dye Backfills

Spinal dye backfills were performed on some lesioned fish after behavioral assays were complete to confirm that vestibulospinal neurons had not regenerated during the behavioral window. To label spinal-projecting neurons in the hindbrain, larvae were anesthetized in 0.2 mg/mL MESAB and mounted laterally in 2% low-melting temperature agarose. Agarose was removed above the spinal cord at the level of the cloaca. An electrochemically sharpened tungsten needle (10130-05, Fine Science Tools, Foster City, CA) was used to create an incision in the skin, muscles, and spinal cord of the larvae. Excess water was removed from the incision site, and crystallized dye (dextran-conjugated Alexa Fluor 546/647 dye (10,000 MW, ThermoFisher Scientific D-22911/D-22914) was applied to the incision site using a tungsten needle. Larvae were left in agarose for at least 5 minutes after dye application before E3 was applied and the fish was removed from agarose. Fish were allowed to recover in E3 for 4-24 hours before imaging. Due to variations in exact incision location and size, spinal dye backfills label fewer spinal projecting neurons than our optical photofill approach and so this method was only used when optical photofills were not feasible.

### Vestibulospinal Photoablations

Vestibulospinal photoablations were performed on *Tg(nefma::EGFP)* larvae at either 4 dpf for use in behavioral experiments from 4-6 dpf, or at 6 or 7 dpf for use in behavioral experiments from 7-9 dpf. Additional experiments were performed on *Tg(α-tubulin:C3PA-GFP)* larvae after spinal photoconversion to target a larger number of vestibulospinal neurons. Larvae were anesthetized in 0.2 mg/ml MESAB and then mounted in 2% low-melting point agarose. Photoablations were performed on an upright microscope (ThorLabs) using a 80 MHz Ti:Sapphire oscillator-based laser at 920 nm for cell visualization (SpectraPhysics MaiTai HP) and a second, high-power pulsed infrared laser for two-photon mediated photoablation (SpectraPhysics Spirit 8W) at 1040 nm (200 kHz repetition rate, 500 pulse picker, 400 fs pulse duration, 4 pulses per neuron over 10 ms) at 25-75 nJ per pulse, depending on tissue depth. Sibling controls were anesthetized for matched durations to lesioned fish. Lesioned and control sibling larvae were allowed to recover for 4-24 hours post-procedure and were confirmed to be swimming spontaneously and responsive to acoustic stimuli before behavioral measurements.

### Behavioral Measurement

Behavioral experiments were performed beginning at either 4 dpf or 7 dpf. For 4 dpf lesions, *Tg(nefma::EGFP)* experiments were performed on 54 vestibulospinal lesioned larvae, and 54 unlesioned sibling controls (5 paired clutch replicates). For 7 dpf lesions, *Tg(nefma::EGFP)* experiments were performed on 97 vestibulospinal lesioned larvae, and 76 unlesioned sibling controls (9 paired clutch replicates). *Tg(α-tubulin:C3PA-GFP)* experiments were performed on 17 vestibulospinal lesioned larvae, and 17 unlesioned sibling controls (5 paired clutch replicates) at 7 dpf. Larvae were filmed in groups of 1-8 siblings in a glass tank (93/G/10 55×55×10 mm, Starna Cells, Inc., Atascadero, CA, USA) filled with 24-26 mL E3 and recorded for 48 hours, with E3 refilled after 24 hours. Experiments were performed in constant darkness.

As described previously^29,41^, video was captured using digital CMOS cameras (Blackfly PGE-23S6M, FLIR Systems, Goleta CA) equipped with close-focusing, manual zoom lenses (18-108 mm Macro Zoom 7000 Lens, Navitar, Inc., Rochester, NY, USA) with f-stop set to 16 to maximize depth of focus. The field-of-view, approximately 2×2 cm, was aligned concentrically with the tank face. A 5W 940nm infrared LED back-light (eBay) was transmitted through an aspheric condenser lens with a diffuser (ACL5040-DG15-B, ThorLabs, NJ). An infrared filter (43-953, Edmund Optics, NJ) was placed in the light path before the imaging lens. Digital video was recorded at 40 Hz with an exposure time of 1 ms. Kinematic data was extracted in real time using the NI-IMAQ vision acquisition environment of LabVIEW (National Instruments Corporation, Austin, TX, USA). Background images were subtracted from live video, intensity thresholding and particle detection were applied, and age-specific exclusion criteria for particle maximum Feret diameter (the greatest distance between two parallel planes restricting the particle) were used to identify larvae in each image. In each frame, the position of the visual center of mass and posture (body orientation in the pitch, or nose-up/down, axis) were collected. Posture was defined as the orientation, relative to horizontal, of the line passing through the visual centroid that minimizes the visual moment of inertia. A larva with posture zero at any given time has its longitudinal axis horizontal, while +90° is nose-up vertical, and -90° is nose-down vertical.

### Behavioral Analysis

Data analysis and modeling were performed using Matlab (MathWorks, Natick, MA, USA). As previously described^29,41^, epochs of consecutively saved frames lasting at least 2.5 sec were incorporated in subsequent analyses if they contained only one larva. Instantaneous differences of body particle centroid position were used to calculate speed. Interbout intervals (IBIs) were calculated from bout onset times (speed positively crossing 5 mm/sec) in epochs containing multiple bouts, and consecutively detected bouts faster than 13.3 Hz were merged into single bouts.

Numerous properties of swim bouts were calculated. The maximum speed of a bout was determined from the largest displacement across two frames during the bout. Trajectory was calculated as the direction of instantaneous movement across those two frames. Displacement across each pair of frames at speeds above 5 mm/sec was summed to find net bout displacement. Bouts with backwards trajectories (*>*90° or *<*-90°) and those with displacements under 0.3 mm were excluded from analysis. Bout duration was calculated by linearly interpolating times crossing 5 mm/s on the rising and falling phases of each bout. Instantaneous bout rate was calculated as the inverse of the IBI duration. Pitch angle distributions were computed using inter-bout pitch, or the mean pitch angle across the duration of an IBI. Pitch probability distributions were calculated using a bin width of 5° (ranging from *±*90°).

A parabolic function was used to fit the relationship between instantaneous bout rate (*y*) in Hz and deviation from preferred posture(*x*) in degrees, based on the formula

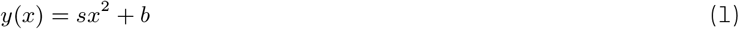

in which *s* gives parabola steepness (pitch sensitivity, in Hz/deg^2^) and *b* gives basal bout rate (Hz). Deviation from preferred posture was itself a function of inter-bout pitch, following the formula:

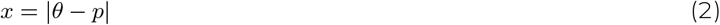

where *θ* is the inter-bout pitch (deg) and *p* is the median inter-bout pitch across all IBIs for each condition. Parameter fits were estimated in Matlab using nonlinear regression-based solver (least-squares estimation). Initial parameter values were *s*=0.001 and *b*=1. Angular velocity sensitivity was calculated by taking the slopes of best-fit lines calculated separately for up and down angular velocities in Matlab using the Theil-Sen estimator. To improve the linear fit, “up” angular velocities were defined as those greater than the median angular velocity and “down” angular velocities were those below the median angular velocity, rather than above/below true zero as this is where the true inflection point of swim rate fell empirically.

Net bout rotation was defined as the difference in pitch angle from 275 ms before to 75 ms after peak speed. Net change in angular velocity was defined as the difference in the mean angular velocity experienced from -225 to - 125 ms (aligned to peak speed) and the mean angular velocity experienced from +250 to +375 ms. For both pitch correction gain and angular velocity gain, parameters were estimated by taking the slope of best-fit line calculated in Matlab using the Theil-Sen estimator.

Total depth change across a bout was calculated as the difference in the vertical position of the fish from 200 ms after peak speed to 250 ms before peak speed. Estimated depth change due to thrust was calculated as the product of the tangent of the fish’s posture at the time of maximum linear acceleration of the swim bout and the empirical horizontal displacement of a bout from -250 to 200 ms aligned to peak speed. Estimated depth change due to lift was calculated ad the difference between total depth change and estimated depth change due to thrust.

### Behavioral Modeling

We generated condition-specific swimming simulations using a generative swim model described previously^29^, with updates to the model to improve the fit to the empirical control dataset collected here. In each condition, 50 simulated fish swam for 3,000 seconds with discrete time steps equivalent to those in the captured data (Δ*t* = 25 ms). Pitch angle (Θ(*t*)) is initialized at a randomly drawn integer from a uniform distribution between *±*90°. At each time step (*t*), pitch angle is updated due to passive posture destabilization from the integral of angular velocity 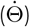 :

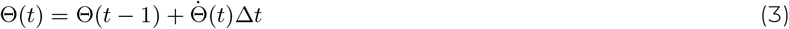

Angular velocity was initialized at 0, and was calculated as the sum of 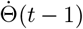 and the integral of angular acceleration, 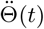 . Between swim bouts, simulated larvae were destabilized according to angular acceleration. Angular acceleration varied as a function of pitch in the preceding time step (Θ(*t* − 1):

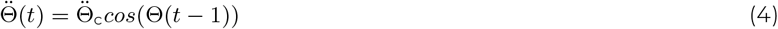

where 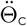 is a maximum angular acceleration for each inter-bout period randomly drawn from a normal distribution centered around 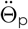 and with a spread of 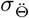, where 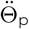 is the the median empirical angular acceleration observed between bouts for fish each condition, and 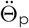 is the inter-quartile range of empirical angular acceleration between bouts.

Simulated larval pitch was updated across time according to passive destabilization until a swim bout was initiated. When a bout was initiated, pitch and angular velocity were updated according to condition-specific correction parameters based on empirical swim bout kinematics. Angular velocity correction from swim bouts was corrected by making net angular acceleration across swim bouts 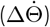 correlated with pre-bout angular velocity 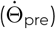. Condition-specific correlations were determined by a single best-fit line to empirical data, defined by a slope 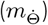 and intercept 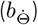. To reproduce the empirical variability of bout kinematics, net angular acceleration incorporated a noise term 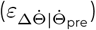 drawn randomly from a Gaussian distribution with mean of 0, and standard deviation 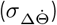 calculated from the empirical standard deviation of 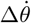 and reduced proportionally by the unexplained variability 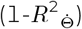 from the correlation between 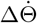 and 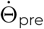.

A bout initiated at time *t* corrected 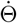 after completion of the bout, 125 ms later (5 time samples, matched to empirical bout duration):

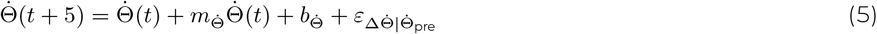

The same approach was used to condition net bout rotation (ΔΘ) on pre-bout pitch (Θ_pre_) based on a single best-fit line, with a single slope (*m*_Θ_) and intercept (*b*_Θ_). For all conditions, the fit line was constrained to the linear portion of the relationship (-30 to +40° pre-bout pitch). If the simulated fish’s posture prior to a bout was outside of this window (6% of simulated control inter-bouts), net bout rotation was calculated using *m*_−30_, *b*_−30_ if Θ *<* −30, or *m*_40_, *b*_40_ if Θ *>* 40. To further impose a ceiling on the counter-rotation values attainable by our simulated larvae, a maximum net bout rotation was imposed (ΔΘ_max_=*±*60°) based on empirical values; if ΔΘ *>* ΔΘ_max_, then ΔΘ was set to ΔΘ_max_. As with angular velocity correction, net bout change also incorporated a noise term drawn from a randomly from a Gaussian distribution with a mean of 0, and a standard deviation ((*σ*_ΔΘ_) calculated from the empirical standard deviation of net bout rotations limited to rotations occurring when pre-bout pitch was between -30 to +40°. Noise was reduced proportionally by the variability unexplained by the correlation (1-*R*^2^_Θ_).

Bouts occurred in the model based an internal state variable representing the probability of bout initiation (*P* _bout_). Bout initiation was calculated as the sum of a non-posture dependent baseline bout rate parameter (*β*), a Θ-dependent bout rate, and -dependent bout rate:

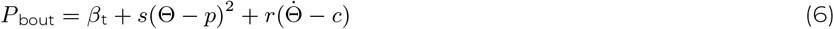

Bout rate as a function of pitch was calculated by fitting a parabola to the relationship between instantaneous bout rate and deviation from mean posture (Θ − *p*) to get a pitch sensitivity parameter (*s*) for each condition. Bout rate as a function of angular velocity was calculated by fitting a line to the correlation between instantaneous bout rate and mean-subtracted angular velocity 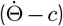 to get a slope (*r*). Instantaneous bout rate increased linearly with the absolute value of 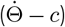 to account for this, two lines were fit, one for negative values of 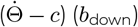 and one for positive values (*b*_up_) to calculate the slopes of the two best-fit lines (*r*_down_, *r*_up_).

Bout initiation probability was updated over time based on Θ(*t*) and 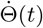, and was also made time-variant, with bout probability dropping to 0 in the first 250 ms (10 time samples) following a bout to match the empirical swim refractory period. After the 250 ms refractory period, swim probability increases to the full bout probability as a linear function of time elapsed in seconds from the last bout (*t*_elapsed_ = *t* − *t_bout_*):

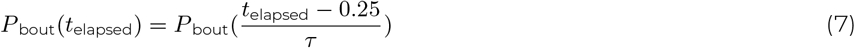

where the rise-time end parameter, *τ* (in seconds), represents the time at which swim probability returns to the full bout probability. The rise time parameter (*τ*) and baseline bout rate parameter (*β*) were fit to minimize the difference between simulated inter-bout interval distribution with the empirical control distribution.

To test that the modified version of swim simulation described here behaved in a similar manner to our previously published model^29^, we compared our full model (described above) using parameters fit from control data with two null models with altered bout initiation and 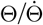 bout correction terms in the model. In the “Bout Timing Null Model,” bouts were initiated in a pitch and angular velocity-independent manner, such that P_bout_ was equal to the the baseline bout rate at all times, but was still time-variant.

In the “Bout Correction Null Model”, Θ_*pre*_ and 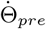 were not correlated with ΔΘ or 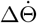, and were instead drawn randomly from a Gaussian distribution with mean and standard deviation matched to empirical ΔΘ or 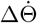 distributions. In the “Complete Null Model” both bout initiation and bout correction terms of the model were randomly drawn. Simulated bouts from the Bout Timing Null Model and Complete Null Model were less well balanced than the Full Model, and performed similarly to our previously published model (Figure S3A). To quantify the discriminability between the simulated pitch distributions of our full and null models and empirical pitch distributions from control larvae, we used the area under the receiver operating characteristic (AUROC). Non-overlapping distributions have an AUROC of 1 and identical distributions have an AUROC of 0.5. The comparison of the Full Control model to the empirical control data had an AUROC of 0.53 at 4 dpf and 0.40 at 7 dpf. The comparison of the Full Lesion model to the empirical lesion data had an AUROC of 0.57 at 4 dpf and 0.47 at 7 dpf.

To test the effects of vestibulospinal neuron lesions on pitch distributions of simulated larvae, we replaced subsets of parameters in the Full Control model with parameters fit from vestibulospinal lesion empirical data. In the Passive model, median posture (*p*), angular velocity (*c*) and angular acceleration 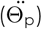 were replaced with parameters fits from lesion data. In the Bout Timing model, *P_bout_*(*t*) was calculated using *β*_*lesion*_, *s*_*lesion*_, *r*_*lesion*_ with all other parameters calculated from control data. In the Bout Correction model,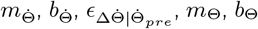, and 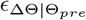 were replaced with parameters fit from lesion data, and all other computations calculated with control parameters.

### Statistics

Due to the large amount of data that we have previously determined to be necessary for behavioral analyses^31^, a single experiment often did not contain enough data to sufficiently sample the entire distribution of postures (3,120*±*1,216 bouts per clutch for 4 dpf experiments; 1,922 *±* 1,416 bouts per clutch for 7 dpf *Tg(nefma::EGFP)* experiments). Therefore data was pooled across all bouts in a given group (lesion or control) when the value being estimated was a fitted parameter or the standard deviation of the distributions, which are largely dependent on the number of bouts included in the analysis. Values reported in the text for fitted parameters and estimates of standard deviation are the estimate across all data per condition. To estimate the variation in parameter fits or in posture standard deviation, we computed 100 bootstrap resampled datasets and calculated the fit/standard deviation on each resampled dataset. Estimates of 95% confidence intervals (shown as error bars in relevant figures) for parameter fits were calculated by finding the values corresponding to 2.5 and 97.5 percentiles of the bootstrapped fits. For bootstrapped parameters, significance testing comparing between conditions was calculated by computing bootstrapped means for each condition and took their differences for statistical analysis. P_bootstrap_ was calculated as a two-tailed significance test, using the mean and standard deviation of differences.

Behavioral statistics for swim kinematics and median posture were calculated as the mean and standard deviation or median and median absolute difference across experimental clutch replicates. When data satisfied criteria of normality, parametric statistical tests (Student’s unpaired t-tests) were used, otherwise we used their non-parametric counterparts (Wilcoxon rank-sum test). Paired statistics were used when comparing the same animals across time (noted).

Expected values and variance of simulated data from computational models were calculated as the mean and standard deviation of mean posture and posture standard deviation across 50 simulated fish. One-Way ANOVAs were used to calculate the effect of changing the parameter sets used in the model on posture standard deviation, and Tukey’s post-hoc tests were used to investigate statistical differences between model versions.

### Data sharing

All raw data and code for analysis are available at the Open Science Framework DOI 10.17605/OSF.IO/GVTHX.

## RESULTS

### Loss of vestibulospinal neurons at 7 dpf disrupts postural stability more strongly than at 4 dpf

Previous electrophysiological^35,37^ and imaging^38^ studies indicate that the larval zebrafish vestibulospinal circuit is poised to regulate posture by encoding body tilt and translation. To determine the specific vestibulospinal contributions to postural stability, we adopted a loss-of-function approach. The transgenic line *Tg(nefma::EGFP)*, reliably labels ∼30 vestibulospinal neurons per brain at both 4 and 7 dpf (60-70% of the total population at both ages, Figure S1A). We could routinely ablate the majority of labelled neurons (22*±*4 neurons at 4 dpf, 23*±*5 neurons at 7 dpf, 50% of the total population at both ages) without damaging surrounding neurons or neuropil (Figure 1A). Photoablated neurons remained absent from larvae 3 days after the lesion occurred suggesting no vestibulospinal recovery or regeneration occurred (1.9*±*2.3 neurons labelled by spinal dye backfill in lesioned hemisphere vs 10.4*±*3.3 in control hemisphere; paired t-test p=5.2×10^-5^).

**Figure 1:**
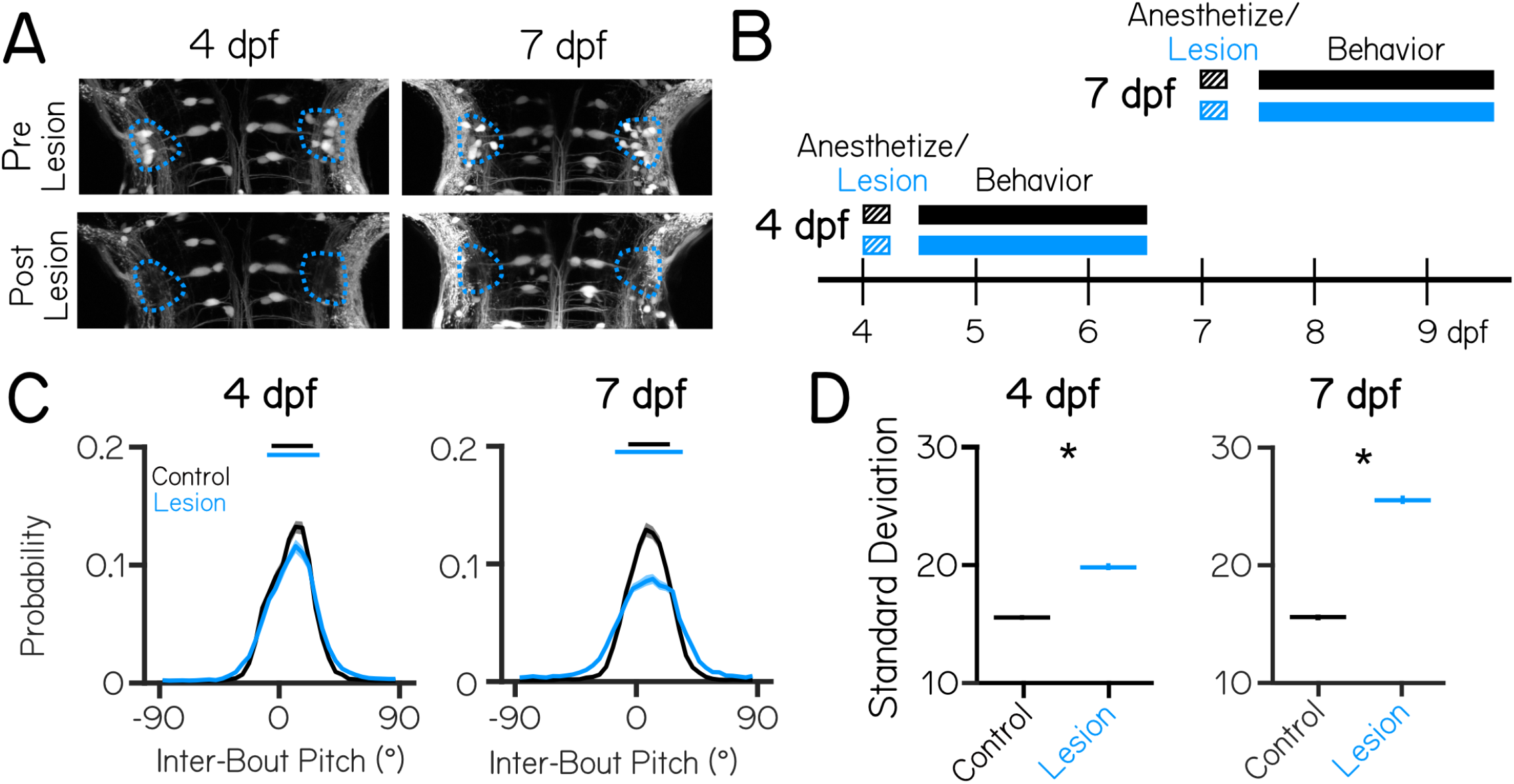
Vestibulospinal neurons contribute to postural stability with increasing effects in older fish. (A) Images taken before and after targeted photoablations of genetically-labeled vestibulospinal neurons show selective loss of fluorescent somata in outlined area (blue) at lesions performed at either 4 dpf (left), or 7 dpf (right). (B) Timeline of the experimental strategy. Fish were either vestibulospinal-lesioned (blue) or non-lesioned sibling controls (black) that were allowed to recover for 4 hours post-lesion and then assayed for postural behavior for 48 hours. Separate experiments were run for fish assayed with lesions at 4 dpf or lesions at 7 dpf. (C) Probability distributions of observed pitch show no change to average posture but greater variability (solid horizontal lines median *±* 1 S.D.) between control fish (black) and lesioned siblings (blue).Solid distribution lines represent probability distribution across all bouts in that condition (4 dpf n=13,548/14,460 bouts Control/Lesion; 7 dpf n=13,382/18,419 bouts Control/Lesion). Shaded error bars represents 95% confidence intervals from 100 bootstrapped resamplings of probability distributions. (D) Postural variability (standard deviation of pitch) is greater in lesioned fish than sibling controls; lines are population standard deviation, error bars represent bootstrapped estimate of 95% confidence intervals. Asterisks represent statistically significant differences, p<0.05.

We used an automated machine vision-based assay^29,31^ to measure locomotion and posture in the pitch axis (nose-up/nose-down) as larvae swam freely in the dark. We measured swim-related kinematics, and postures adopted in between swim bouts. We performed lesions at either 4 or 7 dpf, and monitored locomotion and posture for a 48 hour period after a minimum 4 hour post-procedure recovery period (Figure 1B). Behavior was recorded from either vestibulospinal-lesioned fish, or from sibling controls that were anesthetized for the procedure duration but not lesioned (n = 5 clutch replicates lesions, 54 lesioned fish and 54 lesioned control siblings at 4 dpf; n = 9 clutch replicates, 97 lesioned fish and 76 control siblings at 7 dpf).

We first examined whether loss of vestibulospinal neurons at 4 dpf disrupted the ability of larvae to maintain posture. After vestibulospinal lesions, the distribution of inter-bout pitches adopted by the fish was broader (standard deviation of pitch distribution = 15.6° controls vs 19.8° lesioned, P_bootstrap_=4.8×10^-100^) (Figures 1C and 1D). Lesioned fish were more likely to be found at extreme nose-up or nose-down postures (absolute pitch angle greater than 45°) compared to their siblings (1.3% of inter-bout observations in control vs. 4.3% of lesioned observations) and less likely to be found near (*±*5°) to their preferred posture (25.0% control vs 21.8% lesions). However, the preferred postural set point was not affected (median posture 10.7° controls vs. 11.2° lesions; P_rank sum_=0.84). As the behavioral assay lasted for 48 hours from 4 to 6 dpf, we reasoned that the effect of the lesion might change across this period if the effect of vestibulospinal loss changed within this time window, or if recovery occurred. We found that lesioned fish were more unstable that controls in both the first 24 hour period post-lesion (pitch standard deviation: 16.6 controls vs. 21.7 lesions, P_bootstrap_=1.6×10^-63^, n=5,732/6,428 bouts Control/Lesion) and the second day (14.7 controls vs. 17.5 lesions, P_bootstrap_=9.0×10^-24^, n=7,812/7,995 bouts Control/Lesion). However, in 4 dpf lesions, the magnitude of the effect on the second day post-lesion was smaller than during the first day (standard deviation increased 30% Day 1 vs. 19% Day 2). We conclude that vestibulospinal loss impairs stability across the entire recording period, though some compensation may occur after lesion at 4 dpf. This finding indicates that vestibulospinal neurons contribute functionally towards maintaining stable posture and that this contribution is behaviorally-relevant by 4 dpf.

At 7 dpf, lesioned fish showed more profound disruptions to posture. Lesions at 7 dpf had a broader distribution of inter-bout pitches than their control siblings (15.6° controls vs 25.5° lesions; P_bootstrap_<minval, Figures 1C and 1D). The effect of the lesion on the distribution of observed pitches was stronger at 7 dpf (mean increase in standard deviations 9.9° 7 dpf vs. 4.2° 4 dpf, P_bootstrap_=8.5×10^-80^). While both lesioned and control siblings maintained their posture slightly nose-up from horizontal (median 8.9° control vs. 8.6° lesions), lesioned fish are less likely to be found close to their preferred posture (25.2% control vs 16.9% lesions) and more likely to be found at eccentric nose-up and nose-down postures that control fish rarely adopt (1.4% control vs. 8.5% lesions). Again we observed increased instability in lesions across the 48 hour period of the 7 dpf (standard deviation of pitch distribution: 15.9 controls vs. 25.5 lesions Day 1, P_bootstrap_=5.0×10^-234^, n=6,716/9,790 bouts Control/Lesion; 15.2 controls vs 25.0 lesions Day 2, P_bootstrap_=3.9×10^-231^, n=6,112/8,853 bouts Control/Lesion), though unlike at 4 dpf, the magnitude of the lesion effect on posture standard deviation was consistent across days (38% increase in lesions Day 1 vs 39% increase Day 2) indicating the behavioral compensation did not occur. This result could indicate that developmental timepoint affects the ability to compensate from injury in the vestibulospinal population.

Lesions at either age did not affect basic kinematic properties such as bout speed, bout duration, or distance traveled during a bout (Table 1). Instead, loss of vestibulospinal neurons disrupts the observed distributions of body posture in the pitch axis. Importantly, disruptions to posture are stronger when neurons are lost later in larval development, a key hallmark of a neuronal substrate for balance development.

**Table 1:** Behavioral properties at 4 and 7 dpf.

| Variable | Unit |  |  |  |  |  |  |
| --- | --- | --- | --- | --- | --- | --- | --- |
| Transgenic Line |  | Nefma::GFP | Nefma::GFP | Nefma::GFP | Nefma::GFP | PA-GFP | PA-GFP |
| Condition |  | Control | Lesion | Control | Lesion | Control | Lesion |
| Age | dpf | 4 | 4 | 7 | 7 | 7 | 7 |
| Med. Bout duration (Median ± MAD) | s | 0.13 ($\pm 0.02$) | 0.13 ($\pm 0$) | 0.15 ($\pm 0.01$) | 0.15 ($\pm 0.01$) | 0.13 ($\pm 0$) | 0.13 ($\pm 0$) |
| Med. Bout displacement (Mean ± S.D) | mm | 1.56 ($\pm 0.17$) | 1.47 ($\pm 0.10$) | 1.61 ($\pm 0.18$) | 1.64 ($\pm 0.17$) | 1.53 ($\pm 0.16$) | 1.53 ($\pm 0.16$) |
| Maximum linear speed (Mean ± S.D) | mm/s | 10.8 ($\pm 1.2$) | 10.5 ($\pm 0.8$) | 12.1 ($\pm 1.4$) | 12.1 ($\pm 1.5$) | 11.9 ($\pm 1.6$) | 13.4 ($\pm 1.4$) |

### Vestibulospinal lesions at 7 dpf perturb posture-dependent movement timing and corrective gain computations more than at 4 dpf

Larval zebrafish adopt two behavioral strategies to stabilize posture in the pitch axis. First, translation through the water passively counteracts destabilizing torques^28^. Fish can therefore correct for destabilization by continuously swimming, or by preferentially initiating swim bouts when they sense instability. We previously showed that larvae develop the ability to do the latter^29^ and termed this change to bout initiation “pitch sensitivity.” Second, as part of each bout, larvae actively make angular rotations that partially restore them to their preferred posture^30,31^. These movements are called “corrective rotations,” and their gain (the fraction corrected) increases with age. Together, changes to pitch sensitivity and corrective rotations underlie the development of posture in the pitch axis.

To determine if loss of vestibulospinal neurons interferes with the development of posture, we first assayed pitch sensitivity. The relationship between bout rate and posture is well fit by a parabola (Figure 2A), with two important free parameters: the steepness (pitch sensitivity) and the vertical offset (basal rate of movement). We hypothesized that if vestibulospinal neurons contribute to posture-mediated bout initiation then their loss should (1) flatten this parabola, reflecting decreased pitch sensitivity and (2) increase the basal bout rate to compensate.

**Figure 2:**
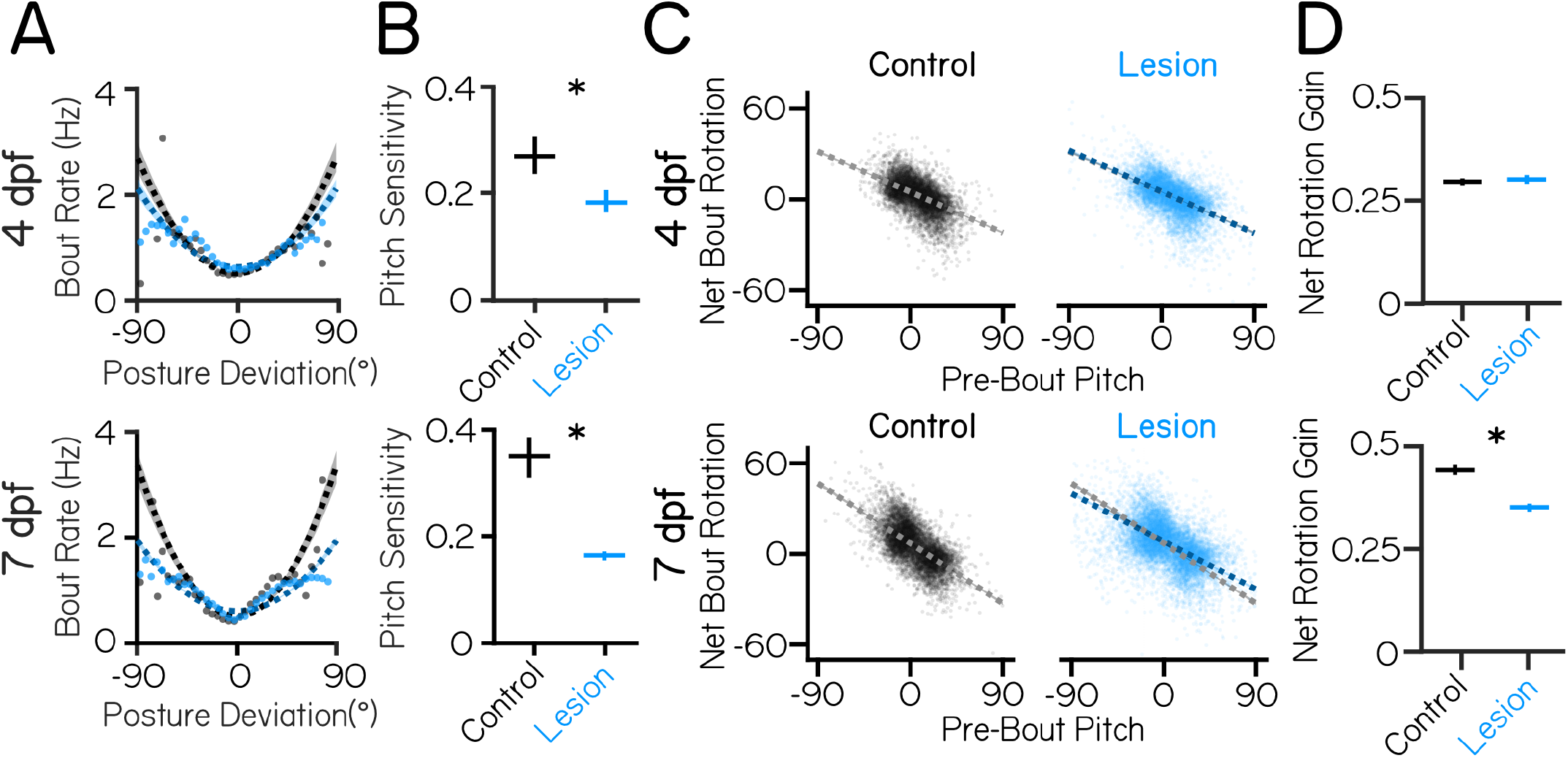
Vestibulospinal neurons contribute to movement timing and corrective capacity with increased effects at 7 dpf than 4 dpf. (A) Movement timing (bout rate as a function of deviation from preferred posture) in lesioned fish (blue) and sibling controls (black). Dots are means of raw data falling within 5° bins. Dashed lines are parabolic fits to raw data (4 dpf n=13548/14460 bouts Control/Lesion; 7 dpf n=13,382/18,419 bouts Control/Lesion). Shaded error bars represents 95% confidence intervals from bootstrapped estimates of fits. (B) Pitch sensitivity (parabolic steepness between posture deviation and bout rate) in lesioned fish and sibling controls at 4 dpf (top) and 7 dpf (bottom). Error bars represent bootstrapped estimate of 95% confidence intervals. (C) Net bout rotation (change in pitch angle before and after a swim bout) as a function of pre-bout pitch in sibling controls (black, left) and vestibulospinal lesioned fish (blue, right). Dots represent individual bouts. Dashed lines are linear fits to included raw data from -30° to +40° Pre-Bout Pitch (4 dpf n=9,979/9,985 bouts Control/Lesion. 7 dpf n=9,238/10,816 bouts Control/Lesion). Control linear fits (gray) are replotted onto lesioned fish data for ease of comparison. (D) Bout rotation gain (absolute slope of the linear fit between pre-bout pitch and net bout rotation) decreases in vestibulospinal lesioned fish at both 4 dpf (top) and 7 dpf (bottom). Error bars represent bootstrapped estimate of 95% confidence intervals. Asterisks represent statistically significant differences, p<0.05.

At 4 dpf, we found that pitch sensitivity was moderately decreased in vestibulospinal lesioned fish (0.27 mHz/deg^2^ controls vs 0.19 mHz/deg^2^ lesioned, P_bootstrap_=5.3×10^-5^) (Figure 2B). Fish lesioned at 7 dpf had a strong decrease in pitch sensitivity relative to control siblings (0.35 mHz/deg^2^ controls vs 0.16 mHz/deg^2^ lesioned, P_bootstrap_=1.6×10^-17^; Figure 2B). Similar to the effects on posture stability, the magnitude of the lesion effect on movement timing computations was greater at 7 dpf (mean decrease in pitch sensitivity=0.08 mHz/deg^2^ 4 dpf vs. 0.19 mHz/deg^2^ 7 dpf, P_bootstrap_=3.9×10^-4^). Consistent with previous findings^29^, we observed that pitch sensitivity was substantially larger in control larvae when measured at 7 dpf than at 4 dpf (0.27 mHz/deg^2^ 4 dpf vs. 0.35 mHz/deg^2^ 7 dpf controls). This developmental increase was entirely absent in lesioned fish assayed at 7 dpf (0.19 mHz/deg^2^ 4 dpf vs. 0.16 mHz/deg^2^ 7 dpf lesioned (Figure 2B). At both timepoints, basal bout rate was significantly higher in lesions compared to control (4dpf: 0.51 Hz controls vs 0.63 Hz lesions, P_bootstrap_=1.3×10^-53^; 7 dpf: 0.50 Hz controls vs 0.59 Hz lesioned, P_bootstrap_=1.9×10^-35^).

As lesioned fish had significantly more observations at extreme pitch angles compared to controls, we reasoned that pitch sensitivity could be lower in lesioned fish due the inclusion of bouts that fall outside of the normal range of observed postures for control fish. To test this, we set a cutoff to exclude the most extreme nose-up and nose-down bouts from the parabolic fits for both control and lesion conditions, and progressively moved to the cutoff from *±*100° from the median (all data included) toward *±*0° (no data included) to determine the cutoff level that removed the effect of lesions on pitch sensitivity. At 4 dpf, the significant effect of lesions on pitch sensitivity disappeared when we excluded bouts more extreme than *±*62.5°, which represents just 1.0% (145 observations) of bouts in the lesion condition and which correspond to postures almost never seen (0.2%, 27 observations) in the control condition. We conclude that at 4 dpf, lesion effects on pitch sensitivity predominately affects only the most extreme postures. At 7 dpf, the significant effect of lesions disappeared when excluding bouts more extreme than *±*37.5°, which represents 11% of the lesion data (2026 observations) and 2% of control data (260 observations). We conclude that changes at 7 dpf are not an artefact of the most extreme posture deviations.

Fish also modulate swim bout frequency as a function of their angular velocity. To test whether angular velocity sensitivity was affected by vestibulospinal lesions, we fit a line to the relationship between bout rate and angular velocity for both negative (nose-down) and positive (nose-up) angular velocity deviations (Figure S2A). Nose-down angular velocity sensitivity was decreased after vestibulospinal lesions at 4 dpf (angular velocity sensitivity nose-down: 0.044 controls vs 0.036 lesion, P_bootstrap_=0.004. Nose-up: 0.061 controls vs 0.061 lesions, P_bootstrap_=0.97)(Figures S2B and S2C). At 7 dpf, lesioned fish showed a weaker relationship between bout rate and both nose-down and nose-up angular velocity prior to a bout (Nose-down: 0.061 controls vs 0.035 lesions, P_bootstrap_=5.3×10^-6^. Nose-up: 0.078 controls vs 0.066 lesions, P_bootstrap_=0.01) (Figures S2A to S2C). Together, these findings suggest that at 4 dpf vestibulospinal neurons contribute modestly towards the increase in movement frequency at unstable pitches/angular velocities but that at 7 dpf, loss of vestibulospinal neurons results in large disruptions in the fish’s ability to preferentially time their swim movements to their experience of instability both at high pitch angle and angular velocities.

We next assessed whether the second key computation that contributes to pitch stability – the ability to generate corrective rotations during a swim bout – was affected by vestibulospinal lesions. Larval zebrafish experience a counterrotation during a swim bout that is negatively correlated with their posture before the swim bout^29,31^; the magnitude of this correlation is the “gain” of the pitch correction. In fish with vestibulospinal lesions at 4 dpf, pitch correction gain was not significantly different from control fish (0.29 controls vs 0.30 lesioned, P_bootstrap_=0.43) (Figures 2C and 2D). In contrast, pitch correction gain was lower in fish lesioned at 7 dpf (0.44 controls vs 0.35 lesions,P_bootstrap_=1.3×10^-27^) (Figures 2C and 2D). Similar to the effects on movement timing, we noticed that pitch correction gain increased in control larvae between 4 and 7 dpf (0.29 at 4 dpf vs. 0.44 at 7 dpf), and that lesions at 7 dpf impaired pitch correction gain such that behavior was similar to that of a younger, intact fish. At both ages lesioned fish performed corrective rotations of comparable magnitudes to those of their control siblings (net rotation 95% intervals [L,U]: 4 dpf controls [-18, 19°], 4 dpf lesions [-22, 21°], 7 dpf controls [-19, 30°], 7 dpf lesions [-23, 34°]), indicating that lesioned fish are capable of making large corrective rotations but do not pair them appropriately to their starting posture. Bouts in lesioned fish also had a small but significant decrease in their capacity to correct for angular velocity instability (Figure S2D) at 7 dpf, supporting the idea that vestibulospinal lesion disrupted the corrective nature of bouts later in development.

While vestibulospinal lesions have clear effects on overall postural stability and the specific computations that help maintain posture, lesioned fish still maintain some level of postural control. To determine whether the residual ability to control posture reflected an incomplete lesion, we repeated our experiments at 7 dpf following optical backfill^43^ of all spinal-projecting neurons in the *Tg(α-tubulin:C3PA-GFP)* line (Figure S1B). Lesions removed the majority of vestibulospinal neurons (n=40*±*8 neurons per fish (83*±*14% of all neurons, n=17 fish), yet produced comparable effects on the standard deviation of pitch distribution (14.0° controls vs 23.3° lesions, P_bootstrap_=1.2×10^-194^) and on pitch sensitivity (0.25 controls vs 0.18 lesions; P_bootstrap_=0.006) as the partial lesions (50% removed) performed in the *Tg(nefma::EGFP)* background at 7 dpf (Figures S1C to S1F). Lesioned fish also had comparably impaired corrective counter-rotations (pitch correction gain: 0.28 controls vs 0.12 lesions, P_bootstrap_=1.9×10^-27^). The remaining ability to maintain posture in these lesions indicates that there are either extra-vestibulospinal contributions to posture, or that a small number of vestibulospinal neurons (8 per fish) can be sufficient to drive an impaired sense pitch sensitivity/corrective counter-rotations. Analysis of 2 lesioned fish that had *>*98% of vestibulospinal neurons removed following optical backfill (0-1 neuron remaining) indicated that while comprehensive lesions did impair stability more strongly than less complete lesions (standard deviation of postures: 26.6 comprehensive lesions, n=2,338 bouts), these fish adopted postures that were still distributed around horizontal (median posture: 1.0 ° comprehensive lesions), suggesting that fish lacking all vestibulospinal neurons are not gravity-blind and maintain some degree of postural control.

Taken together, the results of our lesion experiments support the hypothesis that vestibulospinal neurons play a larger role in postural control as fish develop. We observed that vestibulospinal neurons play a role in postural control behavior as early as 4 dpf. Further, the importance of the vestibulospinal circuit towards postural control increases between the first week of development (4-6 dpf) and early in the second week (7-9 dpf). Specifically, our data argue that loss of vestibulospinal neurons increases variability in the pitch axis by interfering with movement timing at an early developmental age, and posture-dependent movement timing and corrective gain by the second week of larval life.

### A computational model of swimming can explain age-specific consequences of vestibulospinal lesion on postural stability

Loss of vestibulospinal neurons leads to instability in the pitch axis, and disrupts key behaviors that correct posture. Are the changes to disrupted bout timing and/or corrective rotations sufficient to explain the observed instability at both 4 and 7 dpf? To determine the postural impact of vestibulospinal neuron loss, we simulated pitch using a generative model of swimming^29^ (Figure 3A). Model larvae are subject to passive destabilization that is partially corrected by stochastic swim bouts whose kinematics and timing are drawn from distributions that match empirical observations. Of nineteen data-derived parameters (Table 2), thirteen implement the two relevant computations: five that determine the degree to which bout probability depends on either pitch or angular velocity (bout timing) and eight that determine the degree to which bouts restore posture (corrective stabilization). Both bout timing and corrective rotations are required to generate simulated bouts with a preferred horizontal posture and a low-variability pitch distribution (Figure S3A), consistent with previous findings^29^.

**Table 2:** Behavioral modeling parameters.

| Parameter | Symbol | Unit | Control<br>4 dpf | Lesion<br>4 dpf | Control<br>7 dpf | Lesion<br>7 dpf |
| --- | --- | --- | --- | --- | --- | --- |
| Bout duration |  | s | 0.125 | 0.125 | 0.125 | 0.125 |
| Bout refractory period |  | s | 0.25 | 0.25 | 0.25 | 0.25 |
| Median angular acceleration | $\ddot{\theta}_p$ | deg/s<sup>2</sup> | -0.90 | -0.76 | -1.18 | -1.63 |
| Angular acceleration IQR | $\sigma_{\ddot{\theta}}$ | deg/s<sup>2</sup> | 7.6 | 10.1 | 5.8 | 9.0 |
| Median angular velocity | $\dot{c}$ | deg/s | -0.58 | 0.09 | -0.70 | -1.19 |
| Rise-time end | $t_{end}$ | seconds | 1.07 | 1.07 | 1.09 | 1.09 |
| Median posture | $p$ | deg | 10.7 | 11.2 | 8.9 | 8.6 |
| Baseline bout rate | $\beta$ | Hz | 0.36 | 0.36 | 0.30 | 0.30 |
| Pitch sensitivity | $s$ | mHz/deg<sup>2</sup> | 0.27 | 0.19 | 0.35 | 0.16 |
| AV sensitivity (down) | $r_{down}$ | deg<sup>-1</sup> | -0.045 | -0.036 | -0.062 | -0.035 |
| AV sensitivity (up) | $r_{up}$ | deg<sup>-1</sup> | 0.061 | 0.061 | 0.078 | 0.066 |
| Pitch correction gain | $m_{\theta}$ | - | -0.30 | -0.30 | -0.44 | -0.35 |
| Pitch correction intercept | $b_{\theta}$ | deg | 4.9 | 5.1 | 7.0 | 8.2 |
| Net Bout Rotation S.D. | $\sigma_{\Delta\theta}$ | deg | 8.8 | 9.7 | 12.4 | 13.9 |
| Pitch correction R<sup>2</sup> | - | - | 0.30 | 0.27 | 0.40 | 0.27 |
| AV correction gain | $m_{\dot{\theta}}$ | - | -0.80 | -0.83 | -0.93 | -0.87 |
| AV correction intercept | $b_{\dot{\theta}}$ | deg/s | 1.0 | 1.1 | 1.5 | 1.2 |
| Net AV Change S.D. | $\sigma_{\Delta\dot{\theta}}$ | deg/s | 4.9 | 5.0 | 5.3 | 6.2 |
| AV correction R<sup>2</sup> | - | - | 0.51 | 0.45 | 0.50 | 0.39 |

**Figure 3:**
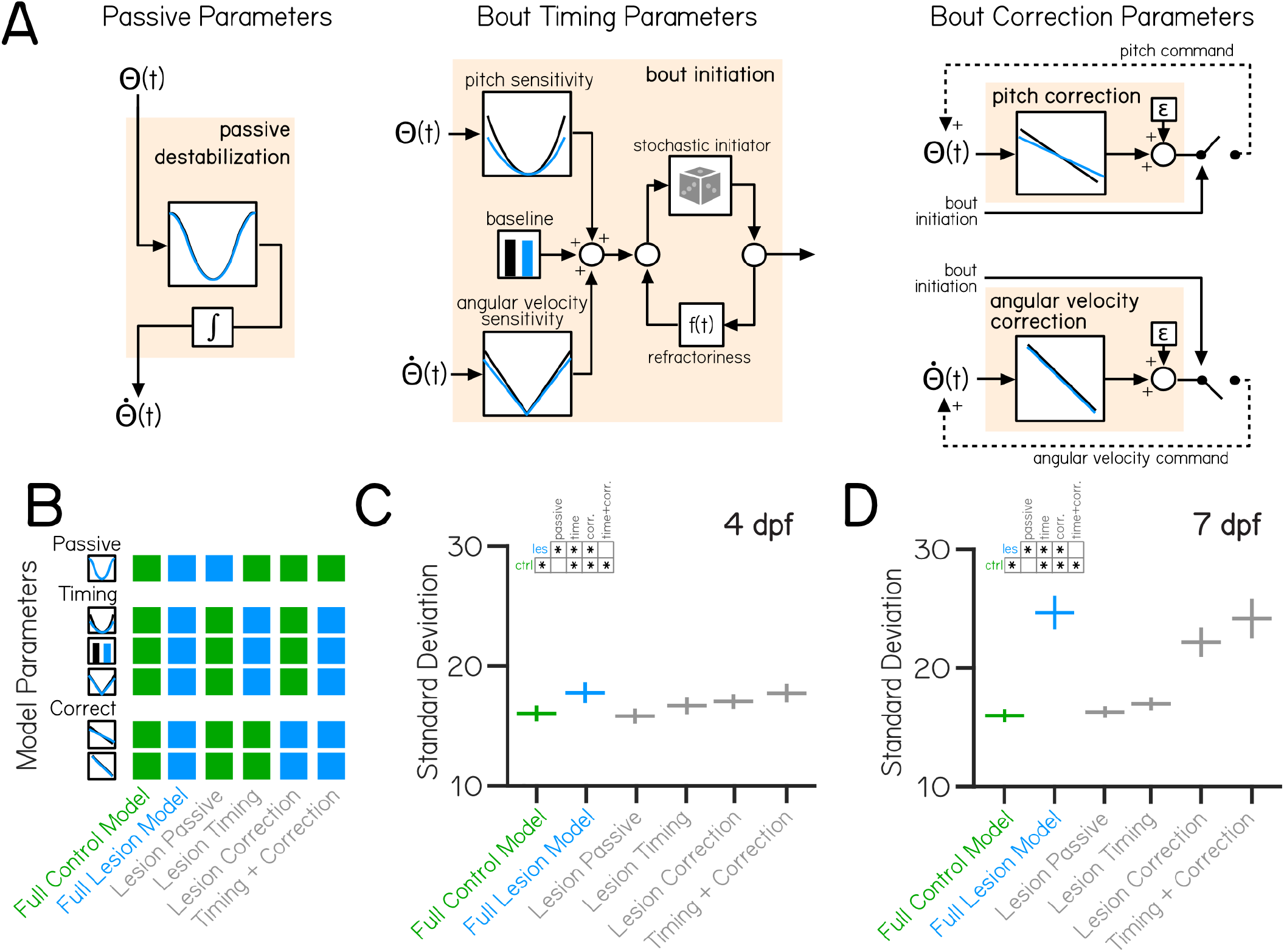
Behavioral modeling shows that increased postural variability following lesions emerges from combined impairments to swim timing and corrective capacity. (A) A generative model of swimming adapted from previously published work^29^, consists of four computations (tan boxes) determining swim timing, pitch angle, and angular velocity. Each computation contains condition-specific parameters, shown by plots with a black (control) or blue (vestibulospinal lesion) line. (B) Variations of the swimming model were created using parameters either entirely from control data (“Full Control Model”, green), lesion data (“Full Lesion Model”, blue), or combinations of parameters from control and lesion data (“Hybrid Models”, gray). (C) Standard deviation of simulated pitch probability distributions *±*S.D. using different combinations of condition-specific parameters based on empirical data from 4 dpf or (D) 7 dpf. Asterisks within inset grids represent statistically significant differences between simulated standard deviations in Tukey’s post-hoc significance tests, p<0.05.

Our model recapitulated the empirical distributions of posture and swim bout timing (Figures S3B to S3D) using parameters derived from control and lesioned data at both 4 and 7 dpf. Modeled pitch distributions had similar mean pitch and standard deviations compared to empirical control data at both 4 dpf (mean posture 12.2*±*1.3° control model, mean distribution standard deviation 16.0*±*0.6° control model) and 7 dpf (mean posture 14.2*±*0.8° control model, mean distribution standard deviation 16.0*±*0.5° control model). Models of control data captured all of the variability (103% of empirical standard deviation) seen in the empirical control distribution at 4 and 7 dpf. Relative to controls, models of lesion data showed broader distributions of pitch angles (mean distribution standard deviation 18.1*±*1.0° lesion 4 dpf model, 24.2*±*1.4° lesion 7 dpf model). Our framework faithfully captures the impact of lesions on posture stability (91% and 94% of empirical standard deviation from 4 and 7 dpf dataset respectively).

If vestibulospinal lesions impair posture stability through disruptions to bout timing and corrective counter-rotations, then changes to both (but not either alone) should explain the increased variability in posture. Alternatively, if one computation alone is sufficient to account for the increased postural variability, then changes in the other computation may be unrelated to effects on stability after lesion. To test how lesion-driven changes to specific computations relate to changes in pitch stability, we systematically replaced the relevant parameters for each computation with those from lesion data (Figure 3B), resulting in a model that is either derived entirely from control parameters (“Full Control Model”, green), entirely from lesion parameters (“Full Lesion Model”, blue) or from a mix of control and lesion parameters (“Hybrid Models”, gray). We then analyzed the standard deviation of the resulting generated pitch distributions from the full control and full lesion models to the hybrid models.

In the 4 dpf hybrid models, the source of parameters used had a signficant effect on the standard deviation of modeled posture distributions (One-Way ANOVA F_5,299_=66.8, p=1.8×10^-46^). Models that replaced only the bout timing parameters with those derived from lesion data had significantly higher standard deviation of postures than the control model (Lesion Timing SD=16.7*±*0.6, p=0.005), but was significantly lower than the full lesion model (p=2.1×10^-8^) indicating that lesion effects on bout timing alone cannot explain the increased instability at 4 dpf (Figure 3C). Surprisingly, though we observed no significant impairment in lesioned fish in parameters controlling the gain of pitch or angular velocity correction at 4 dpf, models that replaced only bout correction parameters also increased in standard deviation compared to controls (Lesion Correction SD=17.1*±*0.7, p=2.1×10^-8^). Only the hybrid model that replaced both bout timing and correction parameters was able to recapitulate the full lesion model (Timing + Correction SD=17.8*±*0.9, comparison to Full Lesion p=0.28). We then further sub-divided the Bout Correction model into additional hybrid models by systematically replacing the eight individual parameters that produce bout correction with the lesion parameters. The lesion parameter contributing the most to the increased standard deviation at 4 dpf was the standard deviation of net bout rotations (*σ*_ΔΘ_) (Figure S3E) which contributes to the amount of variation in the pitch correction computation.

In hybrid models based on 7 dpf data, parameter combinations again significantly impacted the modeled posture standard deviations (One-Way ANOVA F_5,299_=754, p=2.4×10^-165^). Replacing the bout timing parameters (Lesion Timing SD=17.0*±*0.5, p=6.3×10^-5^) or the bout correction parameters (Lesion Correction SD=22.3*±*1.2, p=2.1×10^-8^) resulted in significantly higher pitch standard deviation compared to the control model (Figure 3D). Unlike the 4 dpf model, analysis of individual parameters indicated that effects on the standard deviation in the Lesion Correction model derive from changes both in the standard deviation of net bout rotations (*σ*_ΔΘ_) and pitch correction gain (*m*_Θ_ )(figure S3E) which work together to make pitch correction more variable and less effective. Alone, neither the Lesion Timing nor the Lesion Correction model recapitulated the variability we observed in the full lesion model. Together, the Timing + Correction hybrid model could (Timing + Correction SD=23.9*±*1.4, no statistical difference between Full Lesion and Timing + Correction model, p=0.56). Our models therefore support the hypothesis that changes to timing and bout correction computations work together at both 4 and 7 dpf to drive increased instability in vestibulospinal-lesioned fish. Additionally, modelling supports our conclusion that the contribution of vestibulospinal neurons to balance increases over development.

### Vestibulospinal neurons contribute towards coordination of fin use and body posture

Previous work has established a role for vestibulospinal neurons in paired appendage (limb) movement during postural corrections^25,26^. While zebrafish do not have limbs, they do have pectoral fins, a likely evolutionary precursor of mammalian forelimbs^44^. Pectoral fins in larval zebrafish can generate lift that, when coordinated with trunk rotations, helps fish to climb in the water column. These coordinated movements rely on vestibular sensation and improve over development^41^. We hypothesized that vestibulospinal neurons in the larval fish might contribute towards vestibular-driven fin function, as they do in limb function in mammals.

As zebrafish larvae swim, they can change depth (i.e. swim up/down) in the water column. Depth change can be achieved through thrust-based or lift-based mechanisms^31,41^. When fish swim, they generate thrust that propels them in the direction they are facing – if the fish is at a non-horizontal posture, that thrust will result in a change in depth (Figure 4A, Thrust Δ Depth). Additionally, the pectoral fins can generate a vertical lift force to raise the fish. Pectoral fin loss eliminates all but thrust-based depth changes^41^. We estimate the depth change due to lift during a bout (Figure 4A, Lift Δ Depth) as the difference between the total experienced depth change (Figure 4A, Total Δ Depth) and the change in depth due to thrust (Thrust Δ Depth). The fins’ contribution to depth changes can be described by the lift gain, defined as the slope of the linear relationship between Total Δ Depth and Lift Δ Depth. Lift gain is 1 when the change in depth can be explained entirely by lift, and a near-zero lift gain occurs when depth changes are not correlated with the generated lift (Figure 4A). If vestibulospinal neurons are involved in generating fin-based lift as larvae climb, loss of these neurons should reduce lift gain.

**Figure 4:**
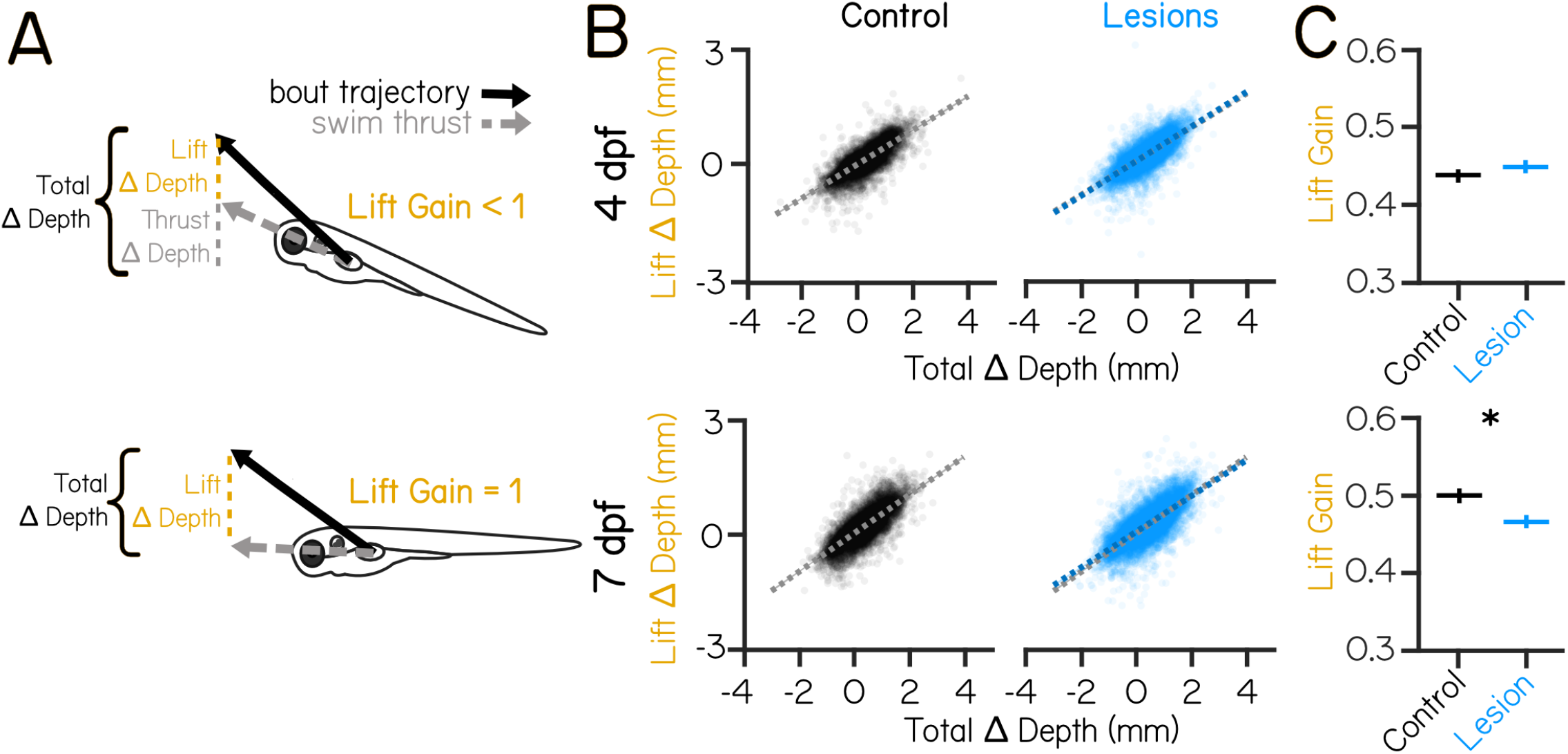
Vestibulospinal lesions disrupt fin and body coordination during vertical navigation only in older larvae. (A) Total depth change that occurs during swim bouts (Total Δ Depth, black) is the sum of depth change due to swim thrust in the direction the fish is pointing (Thrust Δ Depth, gray), and depth change due to vertical lift generated by the fins (Lift Δ Depth, yellow). Lift gain describes the strength of the relationship between Lift Δ Depth and Total Δ Depth. (B) Estimated Lift Δ Depth as a function of the observed Total Δ Depth during a swim bout for control (black, left) and vestibulospinal lesioned (blue, right) fish at 4 dpf (top row) and 7 dpf (bottom row). Dots are single bouts (4 dpf n=10,145/10,454 bouts Control/Lesion; 7 dpf n=9,486/11,663 bouts Control/Lesion). Dashed lines are linear fits to raw data. Shaded error bars represents 95% confidence intervals from bootstrapped estimates of fits. Control linear fits (gray) are replotted onto lesioned fish data for ease of comparison. (C) Lift gain in lesioned fish and sibling controls at 4 dpf (top) and 7 dpf (bottom). Error bars represent bootstrapped estimate of 95% confidence intervals. Asterisks represent statistically significant differences, p<0.05.

Lift gain not significantly changed by vestibulospinal lesions in the *Tg(nefma::EGFP)* line at 4 dpf (0.44 controls vs 0.45 lesions, P_bootstrap_=0.11). In contrast, at 7 dpf, vestibulospinal lesions reduced lift gain (0.50 vs 0.47 lesions, P_bootstrap_=8.0×10^-9^) (Figures 4B and 4C). Lift gain was also significantly decreased in *Tg(α-tubulin:C3PA-GFP)* lesioned fish at 7 dpf (0.36 controls vs 0.30 lesions, P_bootstrap_=7.0×10^-5^). A decrease in lift gain in lesioned fish could arise due to an inability to use the fins to generate lift broadly. However, we observed that the range of all estimated lift magnitudes is comparable between lesioned fish and controls (estimated lift 95% intervals [L,U]: 4 dpf controls [-0.55, 0.78 mm], 4 dpf lesions [-0.56, 0.87 mm], 7 dpf controls [-0.61, 1.04 mm], 7 dpf lesions [-0.60, 1.18 mm]) indicating that lesioned fish can produce fin-based lift of comparable magnitude. A lower lift gain could also result if lift is not coordinated strongly with the overall depth change of the bout. In support of this idea, we saw that lesioned fish had more antagonistic (negative depth change paired with positive lift) swim bouts than control fish at 7 dpf (8.8% of control bouts vs 11.3% of lesion bouts) but not at 4 dpf (8.7% of control bouts vs 7.5% of lesion bouts).

Larval zebrafish with vestibulospinal lesions at 7 dpf, but not 4 dpf, are impaired at coordinating fin-based lift with the appropriate body posture. We conclude that as fish develop, vestibulospinal neurons come to synergize fin and body movements to ensure effective climbs.

## DISCUSSION

Vestibulospinal neurons are an evolutionarily ancient population long thought to play a role in balance regulation. Here we use the larval zebrafish as a model to define the contribution of these neurons to posture control and to understand how that contribution changes over early development. Targeted lesions show that acute loss of vestibulospinal neurons leads to postural instability in the pitch axis. Importantly, this instability is more pronounced in older larvae. Detailed analysis of free-swimming behavior after lesions revealed two failure modes: fish fail to initiate corrective swims appropriately, and their bouts do not adequately restore posture. Once again, the degree to which these balancing behaviors are impaired is age-dependent. *In silico*, it was necessary to incorporate both reduced capacity for corrective restoration and failures of swim initiation to explain variability after lesions at both 4 and 7 dpf. As predicted, larger disruptions in both computations resulted in greater instability in 7 dpf models. Finally, we discovered that at 7 dpf, vestibulospinal neurons contribute to proper fin-body coordination, a key component of vertical navigation that improves with age. Taken together, our data show that loss of vestibulospinal neurons disrupts two key computations – swim initiation and fin-body coordination – that improve between 4 and 7 dpf^29,41^. Vestibulospinal-dependent behaviors therefore play increasingly important roles in postural stability. We conclude that the vestibulospinal nucleus is a locus of balance development in larval zebrafish. Vestibulospinal neurons are found in nearly all vertebrates. We propose that they serve as a partial substrate for a universal challenge: improving posture stability during development.

Our lesion data and model argue that during locomotion vestibulospinal neurons partially facilitate two fundamental computations: specification of the degree of corrective movements and their timing. They do not appear to be necessary for determining postural set point or locomotor kinematics even though larvae can modulate both^29,45^. Similar findings were obtained following partial loss of homologous neurons in the lateral vestibular nucleus, which results in a reduction in the strength of corrective hindlimb reflexes following imposed destabilization^25,26^. Movement initiation (premature stepping) has been observed following stimulation of the lateral vestibular nucleus in cats^46,47^, though see^48^. This previous work was limited to animals that are restrained or performing a balance task; here we advance these studies by demonstrating detrimental impacts to corrective movement timing and gain in naturally-moving animals. Notably, unlike lesions of the vestibular periphery^41,42^,49,50 neither our lesions nor comparable mammalian experiments produce gravity-blind animals, suggesting parallel means of postural control. Similar to observations in the lamprey^10,51^, one parallel pathway likely involves a midbrain nucleus comprised of spinal-projecting neurons called the interstitial nucleus of Cajal, a.k.a. the nucleus of the medial longitudinal fasciculus (INC/nucMLF)^32,45,52–55^. Vestibular information reaches the INC/nucMLF through ascending vestibular neurons in the tangential nucleus^32,42^,54,56. Additionally, it is possible that information about body posture might derive from non-vestibular sensory feedback^57,58^. Taken together, our findings extend complementary loss- and gain-of-function experiments in vertebrates and define one part of the neural substrate for turning sensed imbalance into corrective behaviors.

In addition to regulating computations responsible for postural control, we discovered that loss of vestibulospinal neurons disrupts coordinated fin and body movements zebrafish use to navigate vertically in the water column. Considerable evidence indicates that pectoral fins are evolutionary predecessors to tetrapod forelimbs^44^ that are driven by molecularly-conserved pools of motor neurons capable of terrestrial-like alternating gait^59^. Might vestibulospinal-mediated coordination of trunk and limbs be similarly conserved? Ancient vertebrates without paired appendages such as lampreys have homologous vestibulospinal neurons, but the projections of these neurons terminate in the most rostral portions of the spinal cord and have been postulated to be important in turning but less crucial for the maintenance of posture^10,60^. Vestibular-driven movements of the pectoral fin can be elicited in elasmobranches^61^ and teleosts^32^, indicating that some central vestibular pathway is directly or indirectly connected to the fins. Vestibulospinal neurons in frogs form a key part of the circuit that stabilizes posture at rest and coordinates trunk and hindlimb effectors for balance^62^. In terrestrial vertebrates, postural stability relies on coordination of “anti-gravity” extensor muscles in the trunk and limbs^63–66^. Mammalian vestibulospinal neurons innervate spinal regions that control both ^7–9,33,67–69^. Intriguingly, subsets of vestibulospinal neurons in mice can have functionally different effects on balancing behaviors depending on their downstream spinal cord targets^26^. Evidence in zebrafish also supports the existence of subtypes of vestibulospinal neurons based on sensory afferent input and axon projection type^39^. A key challenge going forward will be identifying transcriptional determinants of subtype identity^70,71^; such a molecular atlas would allow for effective cross-species comparison of subtype function. Our finding that vestibulospinal neurons coordinate fin and trunk movements thus strengthens the proposal that the vestibulospinal circuit serves fundamentally similar roles across disparate body plans and locomotor strategies. By examining the function of vestibulospinal neurons across vertebrate species, we can speculate that vestibulospinal circuits evolved first to maintain posture through trunk effectors and were subsequently adapted to control and coordinate vestibular-drive movement of limbs/limb-like appendages.

As larval fish grow, vestibular-dependent computations involved in posture stabilization and navigation increase in strength^29,30^,41. Here, we identify the vestibulospinal nucleus as a locus for some of these vestibular-dependent computations. We further demonstrate that the vestibulospinal contribution towards these computations increases with age. Specifically, loss of comparable numbers of vestibulospinal neurons had a significantly greater effect on behaviors assayed from 7-9 dpf than at 4-6 dpf. Increases to pitch sensitivity and pitch correction which normally occur between 4 and 7 dpf in non-lesioned fish were partially or entirely prevented by vestibulospinal lesion. We have therefore identified a substrate and a time window during which behaviorally-relevant circuit refinements likely occur. Notably, our findings do not require that physiological changes to the circuit happen within the vestibulospinal neurons themselves. Instead, they implicate changes within this sensorimotor circuit. Additionally, we note that while this study focused only on the effect of lesion at two time-points early in development, behavioral improvement continues until at least 3 weeks post-fertilization^29,41^, meaning that functional refinement of posture circuits is not limited to only the window identified here. Indeed, the postural challenges the fish experience during the first 3 weeks of life increase due to progressive calcification of the anterior skeleton that results in greater passive nose-down angular accelerations^29^. Such an increase in passive instability may demand co-occurring improvements in balance behaviors and the function of underlying neuronal populations. Increased ability to stabilize posture past 9 dpf (the latest timepoint studied here) may be due to increased contribution of vestibulospinal circuits, or to functional changes in other postural control circuits.

Though vestibulospinal neurons were first described over 150 years ago^72^, they remain the focus of active interest to-day^25,26,35,37,57,62,73^. Here we examined the behavioral role of vestibulospinal neurons using precise loss-of-function perturbations with the comparatively simple and well-defined physics of underwater locomotion. We show that vestibulospinal neurons contribute to movement timing, corrective kinematics, and coordination between fin and trunk effectors during navigation. These behaviors are not only fundamental for proper posture and locomotion, but each improves with age^29,41^. The results here indicate that this developmental improvement resides, in part, within the sensorimotor transformation mediated by the vestibulospinal nucleus. Given the near ubiquity of vestibulospinal neurons across vertebrates, our findings are foundational for future studies into the neuronal mechanisms underlying vertebrate postural and locomotor development.

## ACKNOWLEDGMENTS

Research was supported by the National Institute on Deafness and Communication Disorders of the National Institutes of Health under award numbers R01DC017489 and F31DC019554. The authors would like to thank Martha Bagnall along with the members of the Schoppik and Nagel lab for their valuable feedback and discussions.

## AUTHOR CONTRIBUTIONS

Conceptualization: KRH and DS, Methodology: KRH and DS, Investigation: KRH and KH, Resources: YK and SH, Visualization: KRH, Writing: KRH, Editing: DS, Funding Acquisition: KRH and DS, Supervision: DS.

---

**Figure S1:**
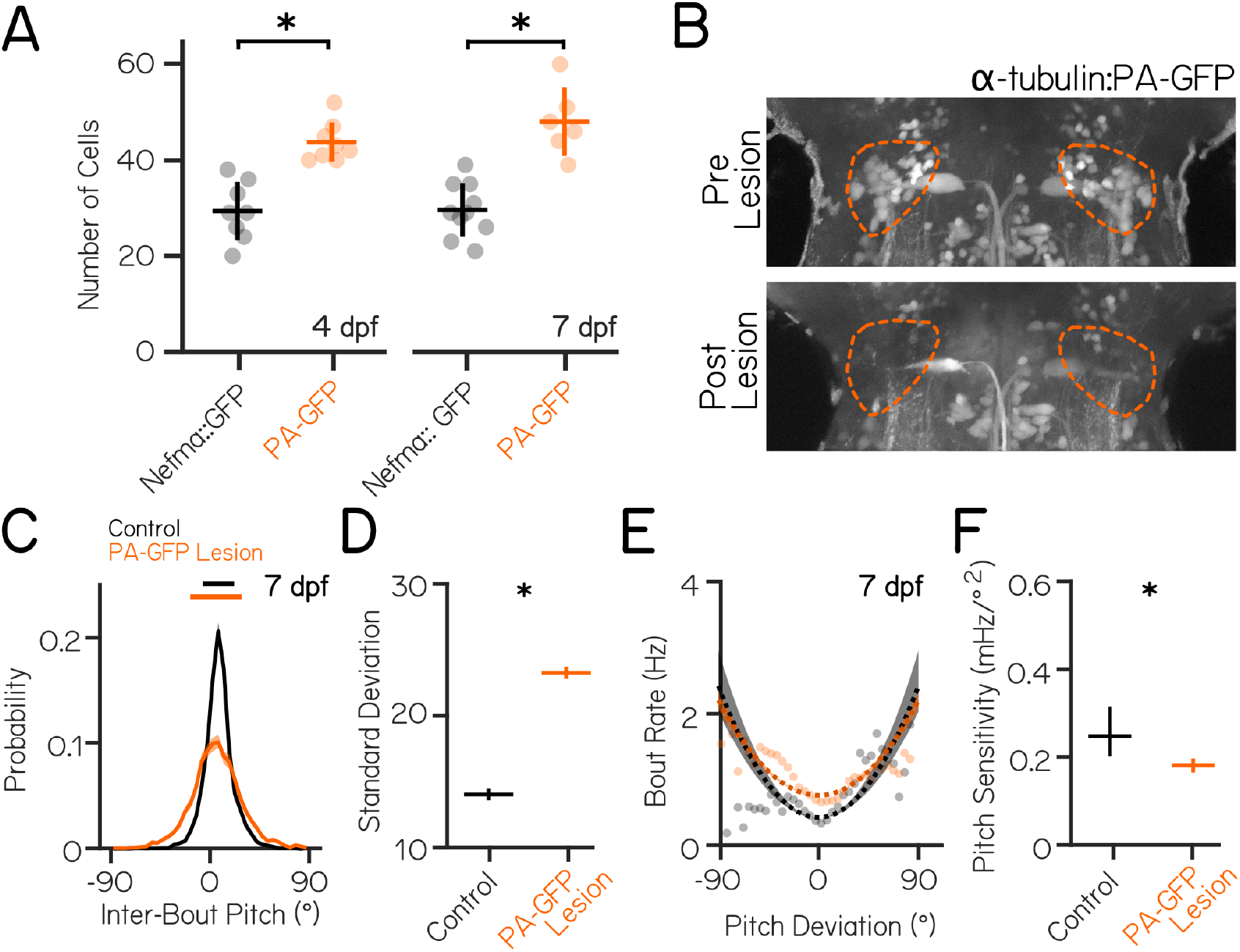
Lesions of a larger pool of vestibulospinal neurons at 7 dpf replicates postural disruption observed in. ***Tg(nefma::EGFP)*** lesions. (A) *Tg(nefma::EGFP)* labels a fraction of the total vestibulospinal population labeled by optically-backfilled *Tg(α-tubulin:C3PA-GFP)* fish when measured at both 4 dpf (29*±*6 cells per fish nefma::EGFP vs. 44*±*4 cells per fish PA-GFP, p=0.0001) and 7 dpf (30*±*6 nefma::EGFP vs. 48*±*7 PA-GFP, p=5.6×10^-6^). Two-Way ANOVA revealed a significant effect of labeling strategy (F_1,31_=63.6, p=1.1×10^-8^), but not of age (F_1,31_=1.2, p=0.29), on number of vestibulospinal cells labeled with no significant interaction effect (F_1,31_=0.96, p=0.34). Dots represent individual fish. Lines represent mean *±* 1 S.D. Asterisks represent statistically significant differences in Tukey’s post-hoc tests, p<0.05. (B) Representative maximum intensity projection of spinal projecting neurons in the hindbrain of a 7 dpf *Tg(α-tubulin:C3PA-GFP)* fish following spinal photoconversions before (top) and after (bottom) two-photon mediated photoablation. (C) Probability distributions of inter-bout pitch angle for sibling controls (black, N=17 fish) and vestibulospinal lesioned fish (orange, N=17 fish) show no change in average posture but greater variability (solid horizontal lines median *±* 1 S.D.) Solid distribution lines represent probability distribution across all bouts in that condition (n=6,510/7,286 bouts Control/Lesion). Shaded error bars represents 95% confidence intervals from bootstrapped estimates of probability distributions. (D) Standard deviation of pitch is higher in vestibulospinal lesioned fish (orange) compared to sibling controls (black). Lines are population standard deviation, error bars represent bootstrapped estimate of 95% confidence intervals. Asterisks represent statistically significant differences, p<0.05. (E) Bout rate as a function of deviation from preferred posture for lesions (orange) and control siblings (black). Solid lines represent raw data, dashed lines represent parabolic fits to raw data, shaded error bars represents 95% confidence intervals from bootstrapped estimates of parabolic fits. (F) Pitch sensitivity (parabolic fit) is decreased in vestibulospinal lesioned fish compared to sibling controls. Error bars represent bootstrapped estimate of 95% confidence intervals.

**Figure S2:**
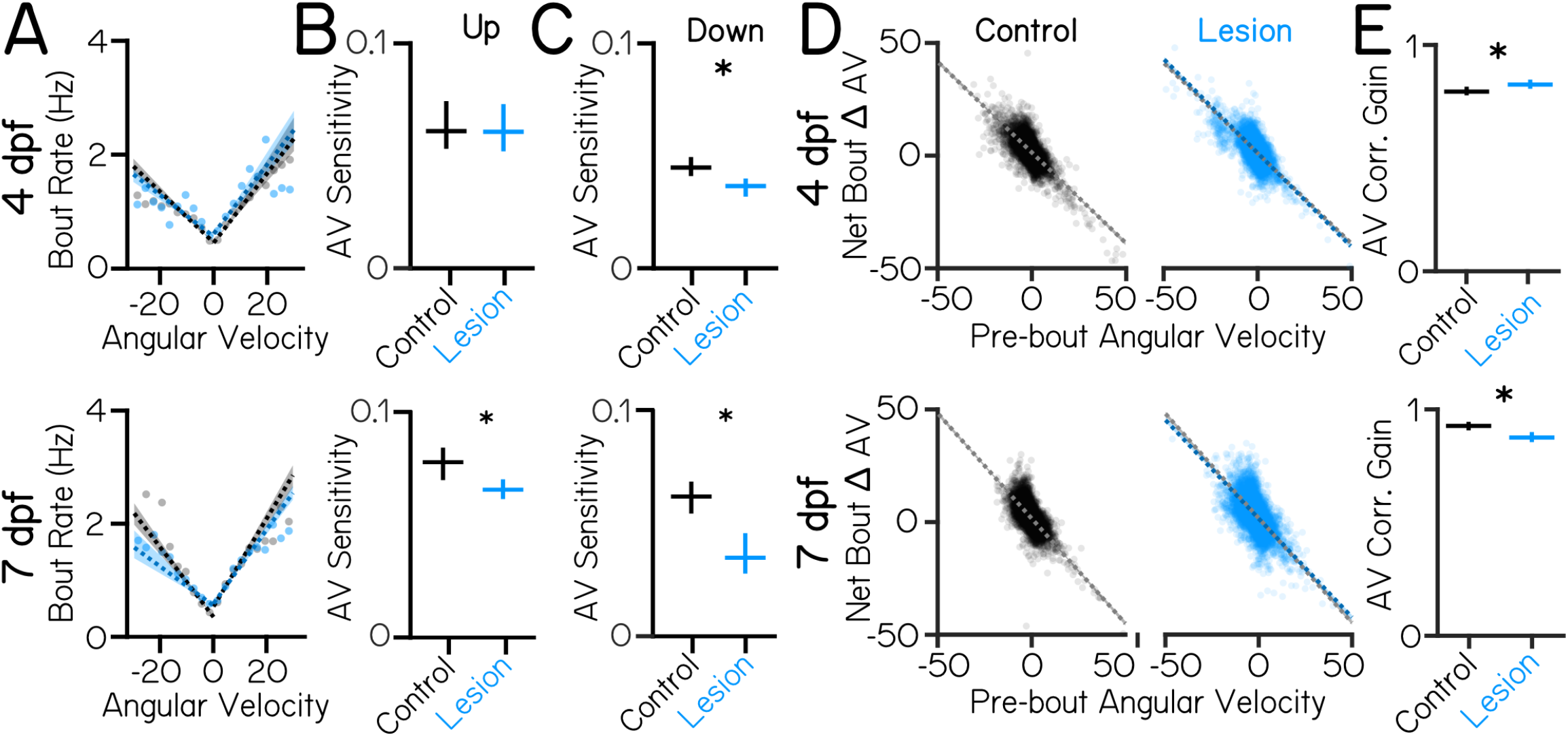
Vestibulospinal lesioned fish have disrupted angular velocity sensitivity and angular velocity correction. (A) Instantaneous bout rate as a function of angular velocity during inter-bout periods in vestibulospinal lesioned (blue) and control siblings (black) at 4 and 7 dpf. Dashed lines represent linear fits to raw data constrained to bouts either below or above the median angular velocity (4 dpf n=6,774/7,320 down bouts, n=6,774/7,230 up bouts Control/Lesion; 7 dpf n=6,691/9,209 down bouts, n=6,691/9,210 up bouts Control/Lesion). Shaded error bars represents 95% confidence intervals from bootstrapped estimates of linear fits. Dots represent binned means of raw data in 3°/s wide bins. (B) Angular velocity (AV) sensitivity (magnitude of linear slope fit) for lesioned and control fish for nose-up and (C) nose-down angular velocities. (D) Net change in angular velocity from the beginning to end of a swim bout as a function of pre-bout angular velocity. Dashed lines represent linear fits to raw data for lesioned (blue, right) or control (black, left) fish. Shaded error bars represents 95% confidence intervals from bootstrapped estimates of linear fits. Dots are individual bouts (4 dpf n=10,225/10,725 bouts Control/Lesion; 7 dpf n=9,625/12,368 bouts Control/Lesion). (E) Angular velocity correction gain (magnitude of linear slope fit) for lesioned and control fish at 4 and 7 dpf. Error bars in panels B, C, and E represent bootstrapped estimate of 95% confidence intervals. Asterisks represent statistically significant effects, p<0.05.

**Figure S3:**
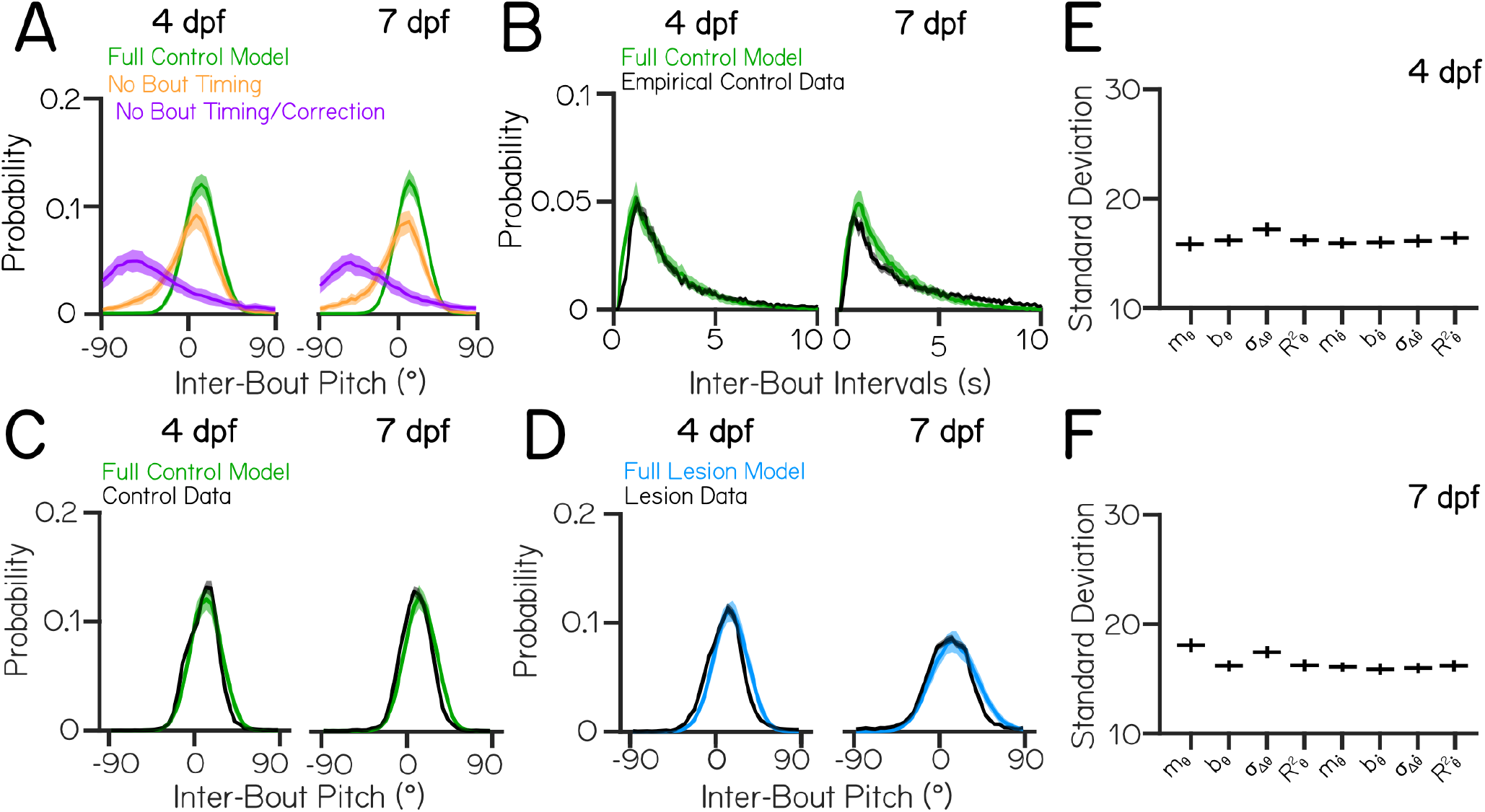
A model of zebrafish swimming during development generates bouts with expected pitch and event-frequency distributions. (A) Probability distributions of simulated inter-bout pitch angles in full control model (green), bout timing null model (purple), and full null model (orange) are comparable for modeling data from 4 and 7 dpf, and are comparable to previous models of larval swimming^29^. (B) Probability distributions of simulated inter-bout intervals for empirical control data (black) and simulated control fish (green) using parameters derived from 4 and 7 dpf sibling control swimming datasets. (C) Probability distributions of observed inter-bout pitch angles at 4 and 7 dpf for empirical control data (black) and simulated control fish (green) and (D) empirical lesion data (black) and simulated lesion fish (blue). For simulated data in panels A-D, lines are probability distribution across all 30 simulated model runs, shaded error bars *±* 1 S.D. across simulated runs. For empirical data in panels A-D, lines are probability distribution across all observations per condition, shaded error bars represent bootstrapped estimates of 95% confidence intervals. (E) Standard deviation of simulated inter-bout pitches in hybrid models with all control parameters except for a single parameter derived from lesion data at 4 dpf or (F) 7 dpf. Parameters shown here calculate pitch bout correction (*m*_Θ_,*b*_Θ_, *σ*_ΔΘ_, *R*^2^_Θ_) and angular velocity bout correction 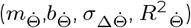 (See Methods, “Behavioral Modeling” for additional explanation of parameters).

